# Immunoglobulin enhancers increase RNA polymerase 2 stalling at somatic hypermutation target sequences

**DOI:** 10.1101/2021.09.16.460442

**Authors:** Alina Tarsalainen, Yaakov Maman, Fei-Long Meng, Minna K. Kyläniemi, Anni Soikkeli, Paulina Budzynska, Jessica J. McDonald, Filip Šenigl, Frederic W. Alt, David G. Schatz, Jukka Alinikula

## Abstract

Somatic hypermutation (SHM) drives the genetic diversity of immunoglobulin (Ig) genes in activated B cells and supports the generation of antibodies with increased affinity for antigen. SHM is targeted to Ig genes by their enhancers (DIVACs; diversification activators), but how the enhancers mediate this activity is unknown. We show using chicken DT40 B cells that highly active DIVACs increase the phosphorylation of RNA polymerase 2 (Pol2) and Pol2 occupancy in the mutating gene with little or no accompanying increase in elongation-competent Pol2 or production of full-length transcripts, indicating accumulation of stalled Pol2. DIVAC has similar effect also in human Ramos Burkitt lymphoma cells. The DIVAC-induced stalling is weakly associated with an increase in the detection of single-stranded DNA bubbles in the mutating target gene. We did not find evidence for antisense transcription, or that DIVAC functions by altering levels of H3K27ac or the histone variant H3.3 in the mutating gene. These findings argue for a connection between Pol2 stalling and *cis*-acting targeting elements in the context of SHM and thus define a mechanistic basis for locus-specific targeting of SHM in the genome. Our results suggest that DIVAC elements render the target gene a suitable platform for AID-mediated mutation without a requirement for increasing transcriptional output.

## INTRODUCTION

Cells work to minimize somatic mutations to avoid genome instability and potential tumorigenesis. Immunoglobulin (Ig) genes are subjected to mutational processes to meet the needs of efficient antigen recognition and appropriate responses of the immune system to pathogenic invaders. Ig gene conversion (GCV) significantly contributes to the primary antibody repertoire of some species, class-switch recombination (CSR) alters antibody effector class, and somatic hypermutation (SHM) provides the variability needed for antigen affinity-based selection of B-cell clones during affinity maturation. These three processes are initiated by the mutator enzyme activation-induced cytidine deaminase (AID) acting on *IgH* switch regions (CSR) or *IgH* and *IgL* V region exons (GCV and SHM) (Arakawa et al., 2002; Muramatsu et al., 1999, 2000; Revy et al., 2000). AID catalyzes the deamination of cytosine in a single-stranded DNA (ssDNA) template, yielding a DNA-resident uracil. Subsequent processing of this lesion follows distinct pathways involving an overlapping set of DNA repair enzymes in CSR, GCV, and SHM (Methot and Di Noia, 2016). In the context of SHM, these lesions are either replicated over or recognized by error-prone mismatch and base-excision repair pathways, eventually resulting in single-nucleotide transition and transversion mutations as well as small insertions and deletions (Methot and Di Noia, 2016).

As AID-induced DNA lesions outside of Ig loci can be detrimental to genomic integrity and can lead to tumorigenesis (Casellas et al., 2016; Robbiani and Nussenzweig, 2013), the activity of AID needs to be tightly regulated. Indeed, the function of AID is regulated at the levels of gene expression, enzymatic activity, nuclear localization, and protein stability among others (Casellas et al., 2016; Xu et al., 2012). AID nuclear localization and target gene transcription alone do not explain the observed mutation window of approx. 150 to 1,500 bp downstream of the Ig transcription start site (TSS) or the preference of SHM for Ig loci, whose mutation frequency is typically orders of magnitudes higher than other transcribed genes in germinal-center (GC) B cells (Casellas et al., 2016; Liu and Schatz, 2009; Liu et al., 2008; Martin et al., 2018). AID can be detected at many regions throughout the genome, even in loci that are not known to undergo somatic hypermutation (Yamane et al., 2011), and recruitment of AID to an IgV region is not sufficient to induce SHM in cells that undergo AID-induced deamination (Matthews et al., 2014). The mechanisms that target AID activity in the genome and, in particular, the mechanisms responsible for the dramatic elevation of AID action at IgV regions relative to other genes remain poorly understood.

AID binds to and travels with RNA polymerase 2 (Pol2), and the source of the ssDNA template for AID is believed to be Pol2-mediated transcription, a strict requirement for SHM (Chaudhuri et al., 2003; Keim et al., 2013; Pham et al., 2019; Yeap and Meng, 2019). Shortly after transcription initiation, DRB sensitivity-inducing factor (DSIF) is recruited to Pol2 together with negative elongation factor (NELF), causing it to pause 25–60 bp downstream of the TSS (Core and Adelman, 2019). This poised Pol2 is phosphorylated at serine 5 of the heptad repeats of the C-terminal domain of the RPB1 subunit (S5P-CTD) by the Cdk7 subunit of the initiation factor TFIIH (Core and Adelman, 2019). Upon pause release, the Cdk9 component of the positive transcription elongation factor P-TEFb phosphorylates serine 2 of the RPB1 CTD (S2P-CTD), NELF, and the Spt5 subunit of the DSIF complex, leading to the dissociation of NELF and allowing the formation of the Pol2 elongation complex (Core and Adelman, 2019). S2P-CTD increases gradually during Pol2 progression through a gene, together with concomitant loss of S5P-CTD (Buratowski, 2009). AID can bind Spt5, and colocalizes with Spt5 and paused Pol2 at S regions (Pavri et al., 2010) as well as at IgV regions in GC B cells (Maul et al., 2014), and associates with components of the Pol2 elongation complex (Methot et al., 2018; Pavri et al., 2010; Willmann et al., 2012).

AID off-target activity leading to chromosomal translocations is associated with antisense transcription and super-enhancers, regions of the genome characterized by a broad and strong pattern of acetylation of histone H3 at lysine 27 (H3K27ac) (Hnisz et al., 2013; Meng et al., 2014; Pefanis et al., 2014; Qian et al., 2014; Whyte et al., 2013). Off-target genes frequently have gene-overlapping inter-connected super-enhancers serving as a source of convergent transcription, which could cause opposing Pol2 collision, stalling of Pol2, and perhaps stabilization of ssDNA on which AID could act (Meng et al., 2014; Qian et al., 2014). AID off-target activity has also been associated with promoter-upstream divergent transcription (Pefanis et al., 2014). These antisense RNA species are degraded by the RNA exosome complex, which may play a direct role in recruitment of AID activity to ssDNA (Basu et al., 2011; Pefanis et al., 2014). Whether antisense transcription and high levels of H3K27ac are required for SHM of IgV regions is not known.

Previous work has provided evidence that targeting of SHM to Ig loci is mediated by *cis*-acting DNA elements that coincide with Ig enhancers and enhancer-like sequences with clusters of TF binding sites (Blagodatski et al., 2009; Buerstedde et al., 2014; Kohler et al., 2012; Kothapalli et al., 2008, 2011; McDonald et al., 2013). These mutation enhancers are collectively called diversification activators (DIVACs) and were identified largely by using SHM reporters. The reporters utilize a GFP expression cassette driven by a strong viral promoter that is largely insensitive to transcriptional enhancement and are integrated into the pseudo V- and Igλ-deleted rearranged *IgL* locus in chicken DT40 B-cell line (Blagodatski et al., 2009; Buerstedde et al., 2014; Kohler et al., 2012; McDonald et al., 2013). When an active SHM recruiting sequence is placed adjacent to the reporter, the GFP gene accumulates AID-induced single-nucleotide substitution mutations, resulting in loss of GFP fluorescence (Blagodatski et al., 2009). DIVACs can increase GFP loss by more than 100-fold without a major effect on transcriptional output (Blagodatski et al., 2009; Kohler et al., 2012). DIVACs rely on NF-κB, IRF/Ets, and MEF2B consensus binding motifs as well as E boxes for full SHM recruiting activity (Buerstedde et al., 2014) and bind E-box binding factors, Ikaros, Ets, and Mef2 family factors in vitro (Dinesh et al., 2020). The endogenous element most clearly associated with the targeting of AID-mediated processes is the 3’RR, a super-enhancer/locus control region, which is critical for efficient CSR and IgH SHM (Cogné et al., 1994; Rouaud et al., 2013; Saintamand et al., 2016; Vincent-Fabert et al., 2010). Previous genome mapping experiments revealed that SHM-susceptibility is a property of certain topologically associating domains (TADs) of chromatin and appears to be confined by TAD boundaries (Senigl et al 2019). Strikingly, when a DIVAC element was inserted into transcriptionally active TADs that do not support SHM, the TADs became SHM susceptible (Senigl et al., 2019). The findings led to a model in which the SHM-susceptibility of non-Ig portions of the genome is driven by non-Ig DIVAC-like elements acting throughout a TAD, perhaps assisted by the chromatin loop-extrusion machinery (Senigl et al., 2019). Similarly, *IgH* locus CSR is driven by chromatin loop extrusion that brings switch regions and the 3′ RR into close proximity (Zhang et al., 2019). These parallels together with the evidence suggesting that Ig and non-Ig super-enhancers can confer susceptibility to AID suggest that the mechanisms that target SHM to Ig loci and to off-target sites in the genome might be closely related. However, the mechanism by which DIVACs stimulate SHM is not known.

To understand how Ig enhancers target SHM, we investigated the effect of DIVACs on constitutively highly transcribed transcription units. We characterized epigenetic marks, a histone isoform previously associated with SHM, antisense transcription, the abundance and phosphorylation status of Pol2, occupancy by Spt5, and single-strandedness at the DIVAC-regulated transcription unit. We observed that DIVAC did not alter many of the parameters that we examined, and that high levels of SHM could occur in the apparent absence of antisense transcription within the mutation target gene. However, we found DIVAC-dependent increases in S2P-CTD and S5P-CTD that co-occur with an increased presence of Spt5 at the transcription unit independently of increased transcript production. In addition, DIVAC can, under certain conditions, increase the single-stranded nature of the mutating target gene, with the increase likely reflecting effects on gene expression as well as mutation load of the target. We conclude that DIVAC modulates RNA polymerase elongation in a way that facilitates AID action on the DNA without necessitating changes in transcriptional output.

## MATERIALS AND METHODS

### GFP reporter constructs

The SD1 DIVAC is a composite DIVAC that consists of human Ig lambda enhancer, chicken light chain enhancer core through 3’ core and human heavy chain intronic enhancer (Buerstedde et al., 2014) (fragments a, b and c in Figure 3A, respectively) and cloned between the *Spe*I and *Nhe*I sites immediately upstream of the pIgL^(-)^ GFP2 plasmid (Blagodatski et al., 2009) to create the *SD1 GFP2* reporter. The *W GFP2* was previously described (Blagodatski et al., 2009). *GFP2* reporters with subregions of W DIVAC, the 1-3, 2-3, 1928 and its E box mutant 1928m upstream were described previously (Kohler et al., 2012; McDonald et al., 2013) To make the *Ri GFP2* reporter, an intronic region from mouse *Rag1* gene (mm10, chr2:101,646,501–101,647,818) was cloned between the *Bam*HI and *Spe*I sites immediately upstream of the RSV promoter of the *GFP2* reporter. The DIVAC 2-3 was cloned downstream of the polyA signal of the *Ri GFP2* between the *Nhe*I and *Bam*HI sites to create the *Ri GFP2 2-3* reporter. The lentiviral *GFP7* and *SD1 GFP7* reporters were described previously but without the T2A sequence (Senigl et al., 2019). To create the *SD2 GFP7* reporter, the mouse Igλ 3-1 shadow enhancer (fragment “d” in Figure 3A) (Buerstedde et al., 2014) immediately 5′to the *SD1 GFP7*.

### GFP loss assay

The GFP loss assay with the GFP2 reporter was done as previously described (Buerstedde et al., 2014). The reporter constructs were transfected to ΔφV IgL^(-)^ AID^R1^ DT40 cells (Blagodatski et al., 2009) and cells with targeted integration to IgL^(-)^ locus selected based on loss of puromycin resistance. A clone with targeted integration was subcloned and cultured for 14 days, after which the percentage of GFP-negative cells in each subclone (minimum of 12) were analyzed with flow cytometry. The GFP loss assay of the *GFP7* reporter was done as described (Senigl et al., 2019). The Ramos cells were transduced with lentivirus at low multiplicity of infection to achieve cells with single integrations. The GFP-positive cells were sorted to isolate single-cell clones. Blasticidin was added after 17 days in culture and minimum of 16 clones were analyzed 21 days after sorting. The GFP-negative percentage of cells in each subclone were analyzed with a flow cytometer. In Figure 1D, each data point represents a median of subclones of an individual experiment as in Figure 1B. A subset of data in Figure 1 B were shown earlier (Kohler et al., 2012; McDonald et al., 2013).

**Figure 1.**
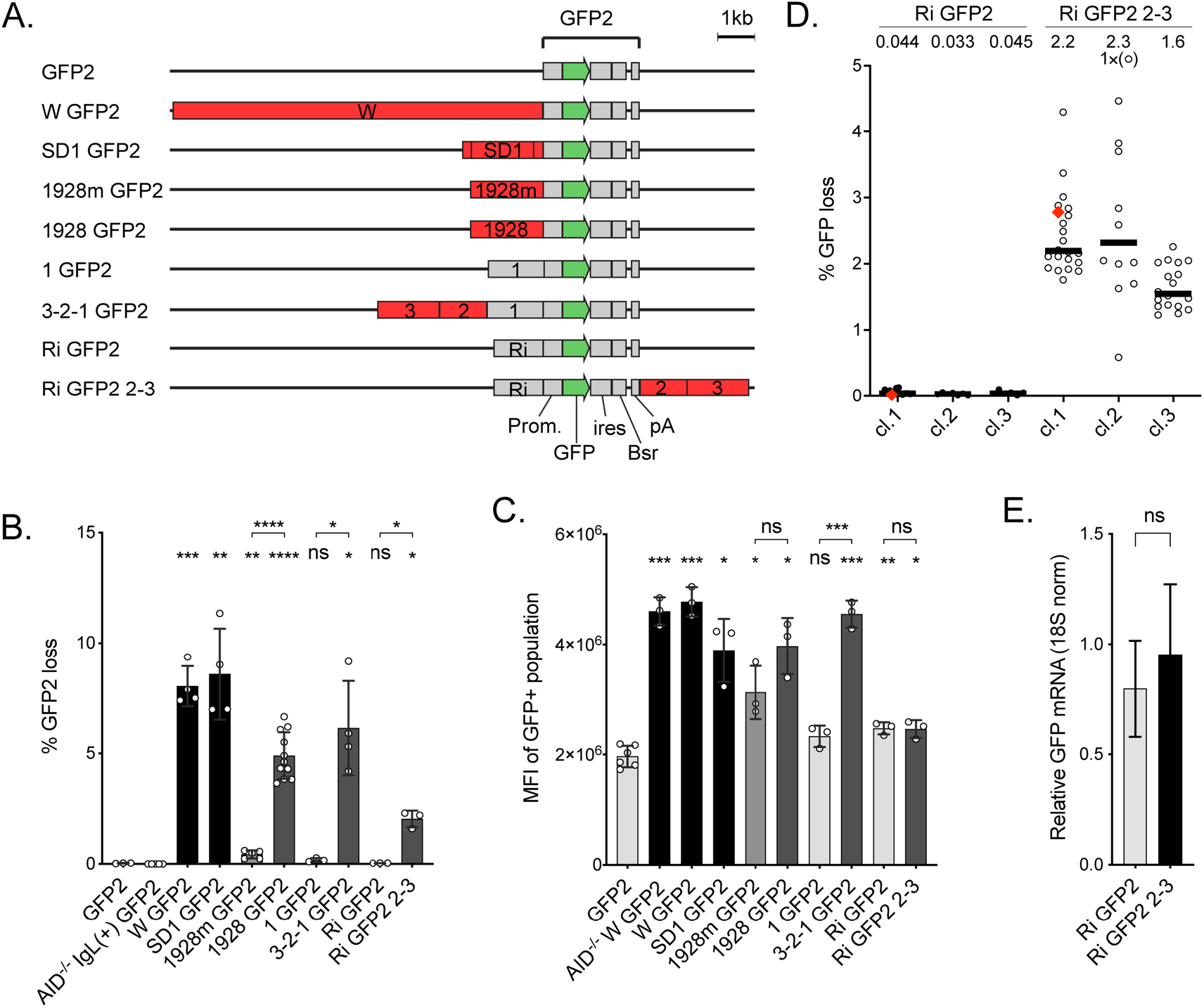
Properties of the SHM reporters used in DT40 cells **A**. Outline of the *GFP2* reporters used in DT40 cells. The reporters differ in the strength (see B) and position of the DIVACs (red) relative to the *GFP* gene (green). The other elements of the reporters are drawn in gray. DIVAC 3-2-1 is a 4.38 kb region of DIVAC W (Blagodatski *et al*., 2009) derived from the chicken *IgL* locus enhancer region and contains subregions 1, 2, and 3 (Kohler *et al*., 2012). DIVAC 2-3 is a 2.99-kb region that lacks subregion 1. DIVAC 1928 is a 1928 bp subregion of 2-3 that was trimmed from the ends and has an internal truncation (Kohler *et al*., 2012). 1928m is E box mutant derivative of 1928 fragment (McDonald *et al*., 2013). Ri is the intronic sequence from the mouse *Rag1* gene. **B**. DIVAC activity (% GFP loss) of the reporters outlined in A. Subregion 1 and Ri do not have DIVAC activity. GFP loss in AID-deficient cell line without a DIVAC or *IgL* deletion is shown as a control (Kohler *et al*., 2012). The bars are shaded to reflect the activities of DIVACs: below 0.2% light gray, 0.2-1% medium gray, 1–6% dark gray, >6% black. Asterisks indicate statistical significance according to Welch’s t-test compared to *GFP2* and relative negative controls as indicated. * *p*<0.05, ** *p*<0.01, *** *p*<0.001, **** *p*<0.0001, ns not significant. **C**. GFP expression in cell lines with targeted integration of the reporters as measured by mean fluorescence intensity (MFI) of GFP-positive cell population. AID-deficiency does not affect the expression of GFP in *W GFP2* reporter (AID^-/-^ W GFP2). Asterisks indicate statistical significance according to Welch’s t-test compared to *GFP2* and relevant negative controls as indicated. * *p*<0.05, ** *p*<0.01, *** *p*<0.001, ns not significant. The shading of the bars is as in B. **D**. GFP loss in three independent clones of DT40 cells with *Ri GFP2* or *Ri GFP2 2-3*, where DIVAC 2-3 is downstream of the *GFP2* cassette. The median percentage of GFP-negative cells after the 14-day assay period of subclones is indicated. One subclone from Ri GFP2 clone 1 and Ri GFP2 2-3 clone 1 each (red diamonds) are used for C, E, and subsequent analyses. Data point outside the y-axis range is in parenthesis. **E**. Expression of GFP mRNA from integrated *Ri GFP2* and *Ri GFP2 2-3* reporters using RT-qPCR. The GFP expression is normalized to 18S RNA. The difference is not significant according to Student’s t-test.

### Expression of GFP

The expression was measured with both GFP fluorescence was analyzed by flow cytometer (NovoCyte, ACEA) and with RT-qPCR. The RT-qPCR RNA was isolated from three cultures using RNeasy Mini kit (Qiagen), the cDNA was synthesized using SensiFAST reverse transcriptase (Bioline), and quantitative PCR was run using SensiFAST Sybr Master Mix (Bioline). The primers used to detect GFP were a1 5′-caagggcgaggagctgttca-3′ and 5′-tgaacttgtggccgtacgtcg-3′ (*GFP7*); a2 5′-cccgaccacatgaagcagca-3′ and 5′-cgctggtggacgtagccttc-3′ (*GFP2* and *GFP7*) and 18S ribosomal RNA were 5′-taaaggaattgacggaaggg-3′ and 5′-tgtcaatcctgtccgtgtc-3′. The signal from GFP was divided by the signal from 18S to normalize.

### Analysis for ssDNA and AID-induced mutations

The sodium bisulfite treatment was performed as described previously (Ronai et al., 2007) with minor modifications. 10^7^ cells were treated with 1.375 M sodium metabisulfite or 2.5 M sodium bisulfite. The DNA was amplified with OneTaq polymerase (New England Biolabs) or with Phusion U polymerase (Thermo Fisher Scientific), cloned with Zero Blunt TOPO PCR Cloning Kit (Invitrogen), and sequenced using Sanger sequencing.

Regions with at least two consecutive converted Cs on the same strand were considered ssDNA patches. SsDNA character was determined by two parameters: the frequency of single-stranded deoxycytidines (ssDNA frequency; number of Cs in patches divided by total number of sequenced Cs) and ssDNA patch size. Patch size is the mean of the minimum and maximum possible sizes of the ssDNA patch. The minimum size is the number of nucleotides from the first to the last converted C, and the maximum size is the number of nucleotides between the first and the last non-converted C. For comparison of constructs, these parameters were assessed within the *GFP* open reading frame (TSS +15 to +714) since it is in this region where mutations can cause GFP loss (Figure 6A-C). For *keratin 5*, 6 sequences of 675 bp were analyzed. The region from TSS +15 to +1,864 (GFP-ires-Bsr) was analyzed for the location of ssDNA patches and the location of AID-induced mutations (Figure 6 E-G).

The GC content was calculated with GC content calculator using the formula Count(G + C)/Count(A + T + G + C) × 100% and plotted with a 30-bp window (Biologics International corp. https://www.biologicscorp.com/tools/GCContent).

### ChIP-seq

The ChIPs were done essentially as previously described (Williams et al., 2016), with the following modifications. 4 × 10^7^ formaldehyde-fixed cells were lysed in SDS lysis buffer (50 mM Tris-HCl, pH 8.0, 10 mM EDTA, 1% SDS, supplemented with Complete, EDTA-free Protease Inhibitor Cocktail (Roche)) and diluted 4-fold with dilution buffer (16.4 mM Tris-HCl, pH 8.0, 167 mM NaCl, 1.2 mM EDTA, 0.01% SDS, 1.1% Triton X-100, supplemented with inhibitors). The lysate was sonicated in aliquots using a water bath sonicator (Diagenode Bioruptor Pico) for 11 cycles (30 s on/30 s off) and cleared by centrifugation. The combined sonicated material was further diluted 1.5-fold with dilution buffer, and 1.7 × 10^7^ cell equivalents were subjected to immunoprecipitation with 5 μg (α-H3K27Ac, α-H3.3, α-RPB1, α-S5P CTD, and α-S2P CTD), 7 μg (α-Spt5), or a 1:50 dilution (α-H3K36me3) of antibodies overnight. The next day, immune complexes were incubated with magnetic beads for 2 h (Protein G Dynabeads, Thermo Fisher Scientific or Protein A/G magnetic beads, Pierce). The beads were collected using a magnetic rack and washed twice with low-salt wash buffer (20 mM Tris-HCl, pH 8.0, 150 mM NaCl, 2 mM EDTA, 1% Triton X-100, 0.1% SDS, supplemented with inhibitors), twice with ChIP wash buffer (50 mM HEPES, pH 7.6, 500 mM LiCl, 1 mM EDTA, 0.7% sodium deoxycholate, and 1% Igepal CA-630), and twice with TE buffer with 50 mM NaCl. Chromatin was eluted in 100 µl of SDS lysis buffer with incubation at 1200 rpm on a thermal mixer for 15 min at 65 °C. The mixture was supplemented with 1 volume of TE and Proteinase K and incubated at 65 °C overnight. The DNA was purified using phenol-chloroform extraction. The following antibodies were used for ChIP: α-H3K27Ac (ab4729), α-S5P CTD (ab5131), and α-S2P CTD (ab5095) from Abcam; α-H3K36me3 (D5A7, #4909) from Cell Signaling Technology; α-H3.3 (09-838) from Millipore; α-RPB1 (N20x) and α-Spt5 (H-300x) from Santa Cruz Biotechnology.

ChIP-seq libraries were prepared for each ChIP from three independent ChIP experiments using TruSeq ChIP Sample Preparation Kit (Illumina) or NEBNext Ultra II DNA Library Prep Kit (New England Biolabs) according to manufacturer’s instructions, but with fewer and ChIP-specific amplification cycles. The libraries were sequenced using the NextSeq 500 instrument (Illumina). The reads were aligned to the relevant custom genomes using Bowtie2 (Langmead and Salzberg, 2012). The genomes used were the Gallus_gallus-4.0 genome, with the region spanning φV and *IgL* locus (chr15:7,921,694–7,955,323) replaced with respective reporter construct sequence and the human GRCh38 genome with the lentivirus reporter appended as an extra chromosome.

The ChIP signals (rpkm) from the GFP transcription unit of the reporter were compared to the 500 most highly expressed genes longer than 1,500 bp (according to DT40 GRO-seq and Ramos GRO-seq (Meng et al., 2014)). Chicken and human gene annotations were from Gallus_gallus-4.0 and GRCh38 genomes, respectively.

### GRO-seq

The GRO-seq was done as previously described (Core et al., 2008). Nuclei were prepared from 20 million cells of each line. The reads from libraries were aligned with Bowtie2 (Langmead and Salzberg, 2012) to the custom Gallus_gallus-4.0 genome with the region spanning φV and *IgL* locus (chr15:7,921,694–7,955,323) replaced with respective reporter construct sequence. The alignment was done as previously described (Meng et al., 2014).

#### Data availability

ChIP-seq and GRO-seq data are available through GEO database (Accession number: GSE180178).

## RESULTS

### Experimental system for analysis of DIVAC function

To investigate the mutation targeting mechanism of DIVACs, we compared GFP-based genome-integrated transcription units either flanked or not flanked by a strong DIVAC. In these systems, the DIVACs can stimulate AID-mediated mutation of the transcription unit up to two orders of magnitude while the production of full-length transcripts and GFP protein are either unaltered or only slightly increased. Two different reporter systems were used, both of which utilize a strong viral promoter driving GFP transcription, downstream of which are an internal ribosome entry site (ires) and a drug selection marker (Bsr). In the first system, the *GFP2* reporter vector (Figure 1A) is integrated into the chicken DT40 B-cell line by homologous recombination, with insertion into an “empty” *IgL* locus lacking IgL and pseudo V sequences (Blagodatski et al., 2009). DT40 cells have been used extensively to characterize Ig GCV, SHM, and AID targeting (Arakawa and Buerstedde, 2009; Arakawa et al., 2002; Blagodatski et al., 2009; Budzyńska et al., 2017; Buerstedde et al., 2014; Chandra et al., 2015; Kodgire et al., 2013; Kohler et al., 2012; Kothapalli et al., 2008, 2011; McDonald et al., 2013; Romanello et al., 2016; Tanaka et al., 2010; Williams et al., 2016; Yabuki et al., 2009). The second system utilizes a retroviral reporter vector and the human Burkitt lymphoma line Ramos and is described below.

After integration of *GFP2*, with or without a flanking DIVAC element, into the DT40 *IgL* locus, single-cell subclones were expanded for about 3 weeks and SHM was measured by the loss of GFP fluorescence (hereafter, “GFP loss”) by flow cytometry. As reported previously (Blagodatski et al., 2009; Kohler et al., 2012; McDonald et al., 2013), *GFP2* alone (“no DIVAC”) yields extremely low levels of GFP loss (median of 0.034%), and *GFP2* in an intact *IgL* locus but in AID-deficient DT40 cells (AID^-/-^ IgL(+) GFP2) yielded even lower levels of GFP loss (0.0038%) (Figure 1B). Several different DIVAC elements were examined (Figure 1A). The ∼10-kb W DIVAC fragment, which contains all known chicken *IgL* DIVAC function (Blagodatski et al., 2009), induces SHM activity 200-fold over the no-DIVAC control (Figure 1 B). DIVAC 3-2-1 is a portion of the W fragment, with DIVAC function nearly as strong as that of W, and consists of three subfragments (1, 2, and 3), of which fragments 2 and 3 contain the DIVAC activity (Figure 1 A and B; compare *3-2-1 GFP2* to *1 GFP2*) (Kohler et al., 2012). Fragments 2 and 3 were further trimmed to a 1,928-bp DIVAC element that retains most of the DIVAC activity of the W fragment (Figure 1A and B; 1928 GFP2) (Kohler et al., 2012). Mutation of all 20 E boxes within DIVAC 1928 (<u>C</u>ANNTG to <u>A</u>ANNTG) reduces its DIVAC activity 10-fold (Figure 1A and B; *1928m GFP2*) (Kohler et al., 2012; McDonald et al., 2013). SuperDIVAC1 (SD1) is a composite 2.1-kb DIVAC element composed of the active core of the W fragment flanked by the human *IgH* intronic enhancer and human *IgL* enhancer core elements (Williams et al., 2016). SD1 drives GFP loss as efficiently as the entire W fragment (Figure 1B; *SD1 GFP2*).

DIVACs have a modest potential to increase the expression of the neighboring transcription unit in the context of genome-integrated SHM reporters. The most potent DIVACs (W, SD1, 3-2-1, and 1928) increase GFP expression by 2–2.5-fold, as assessed by GFP mean fluorescence intensity (MFI) of the GFP-positive population (Figure 1C), which is far less than the stimulation of GFP loss (Figure 1B). AID has no discernable effect on GFP MFI, since the W GFP2 reporter in an AID-deficient cell line had similar GFP MFI in both AID-deficient and in AID-expressing cells (Figure 1C; compare *W GFP2* to *AID*^*-/-*^ *W GFP2*). In the reporters described above, the DIVACs are inserted immediately upstream of the *GFP2* transcription unit and thus might increase GFP expression by providing incremental promoter activity.

To test this idea and to attempt to eliminate the stimulatory effect of DIVAC on transcription, we inserted DIVAC regions 2-3 downstream of *GFP2* (Figure 1A). We also inserted a mouse Rag1 intron (Ri) sequence immediately upstream of the promoter to provide a unique base-balanced sequence to facilitate subsequent analyses. The Ri sequence had no effect on GFP loss (Figure 1D, *GFP2* 0.034% vs. *Ri GFP2* 0.041, *p*=0.50) or on levels of full-length *GFP* transcripts (Figure 1E, *GFP2* 0.80 vs. *Ri GFP2* 0.95, *p*=0.43). Importantly, DIVAC 2-3 did not significantly alter expression from the downstream location, as assessed by GFP MFI (Figure 1C, 3-2-1 GFP2 vs. Ri GFP2 2-3). Three independent clones of both *Ri GFP2* and *Ri GFP2 2-3* were analyzed for GFP loss. DIVAC 2-3 increased GFP loss on average 20-fold relative to the *Ri GFP2* control (Figure 1B, GFP loss *Ri GFP2 2-3* 2.03%, *p*=0.012), indicating that the DIVAC 2-3 has strong DIVAC activity from both upstream and downstream locations. Single subclones from Ri GFP2 cl.1 and Ri GFP2 2-3 cl.1 (Figure 1D, red diamond) that have a non-significant difference GFP MFI (Figure 1C) or GFP mRNA expression as assessed by RT-qPCR (Figure 1E) were used for subsequent analyses. The downstream configuration isolates DIVAC activity from effects on steady-state transcription levels and allows for a well-controlled investigation into the DIVAC targeting mechanism.

### DIVACs do not act through increasing H3K27ac

As noted above, super-enhancers are sometimes themselves targeted by off-target SHM. To understand the contribution of the H3K27ac, a histone mark for enhancers, to the targeting of SHM, we asked whether DIVACs alter levels of H3K27ac in the SHM reporter. ChIP-seq revealed substantial H3K27ac across the entire *Ri GFP2* reporter (Figure 2A). As expected, the DIVAC 2-3 region displayed high levels of H3K27ac and was associated with a small but detectable increase of H3K27ac in the reporter area flanking the DIVAC and the genomic region downstream of the reporter (Figure 2A, panels i and ii).

**Figure 2.**
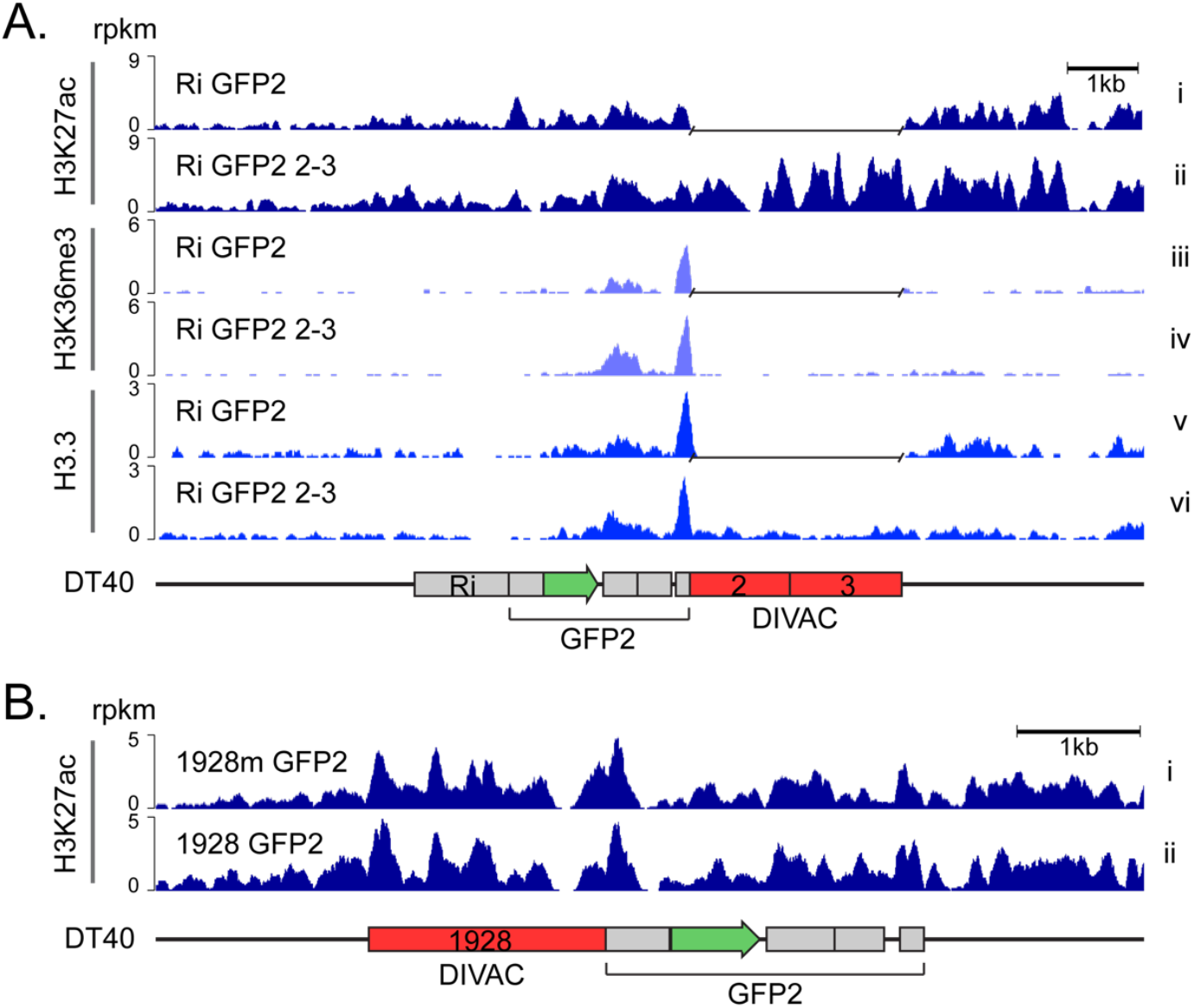
Effect of DIVAC on histone marks in DT40 cells **A**. H3K27ac, H3K36me3 and H3.3 ChIP-seq tracks of genome-integrated *Ri GFP2* (upper tracks) and *Ri GFP2 2-3* (lower tracks) reporters in IgL^(-)^ DT40 cells. A gap has been inserted in the tracks of *Ri GFP2* reporter in the place of the DIVAC 2-3 sequence after the alignment. Values are reads per kilobase per million reads (rpkm). **B**. H3K27ac ChIP-seq of *1928 GFP2* and E-box mutant *1928m GFP2* reporter as in A.

To investigate whether the increase in H3K27ac over the entire reporter was simply caused by the inserted DIVAC 2-3 element, we extended the analysis to *DIVAC 1928* and *DIVAC 1928m*, which differ only by 20 single-bp changes. The H3K27ac pattern and levels in the two reporters were nearly indistinguishable (Figure 2B), indicating that modulation of H3K27ac is unlikely a critical part of the mechanism by which DIVAC stimulates SHM in DT40 cells.

To verify that DIVAC does not act by modulating levels of H3K27ac and that these findings were not a cell line- or species-specific phenomenon, we tested the effect of DIVAC on H3K27ac in the human Burkitt lymphoma cell line Ramos. To this end, we used lentivirus-based reporter for SHM, *GFP7*, in which the GFPnovo2 (Buerstedde et al., 2014) coding sequence is fused in frame to an upstream hypermutation targeting sequence (HTS) (Figure 3A). The HTS contains a high density of AID hotspots, and its sequence was designed such that cytidine deamination frequently leads to in-frame stop codons and the loss of green fluorescence of the reporter-bearing cells. After proviral integration into the genome, transcription of *GFP7* is driven solely by the internal CMV promoter, because the vector contains a self-inactivating 3’ long terminal repeat (Senigl et al., 2019). In the *GFP7* system, as with *GFP2* in DT40, GFP loss is strongly stimulated by DIVACs and is caused by AID-dependent point mutations in the reporter open reading frame (Figure 3A and (Senigl et al., 2019)). The DIVAC elements used in the *GFP7* system were SD1 and a slightly longer version, SD2, which differs by addition of the mouse Igλ 3-1 shadow enhancer (Buerstedde et al., 2014) upstream of the SD1 (fragment “d” in Figure 3A). The lentiviral *GFP7* reporter with SD1 has a more than 30-fold increased mutation rate compared to the no-DIVAC control, as measured by GFP fluorescence loss, and a less than 2-fold induction of transcription (Figure 3B and C). SD2 induces similar or stronger GFP loss (Figure 3B) and a 2-fold increased GFP protein expression level compared to no DIVAC, similar to that seen with SD1 (Figure 3D).

**Figure 3.**
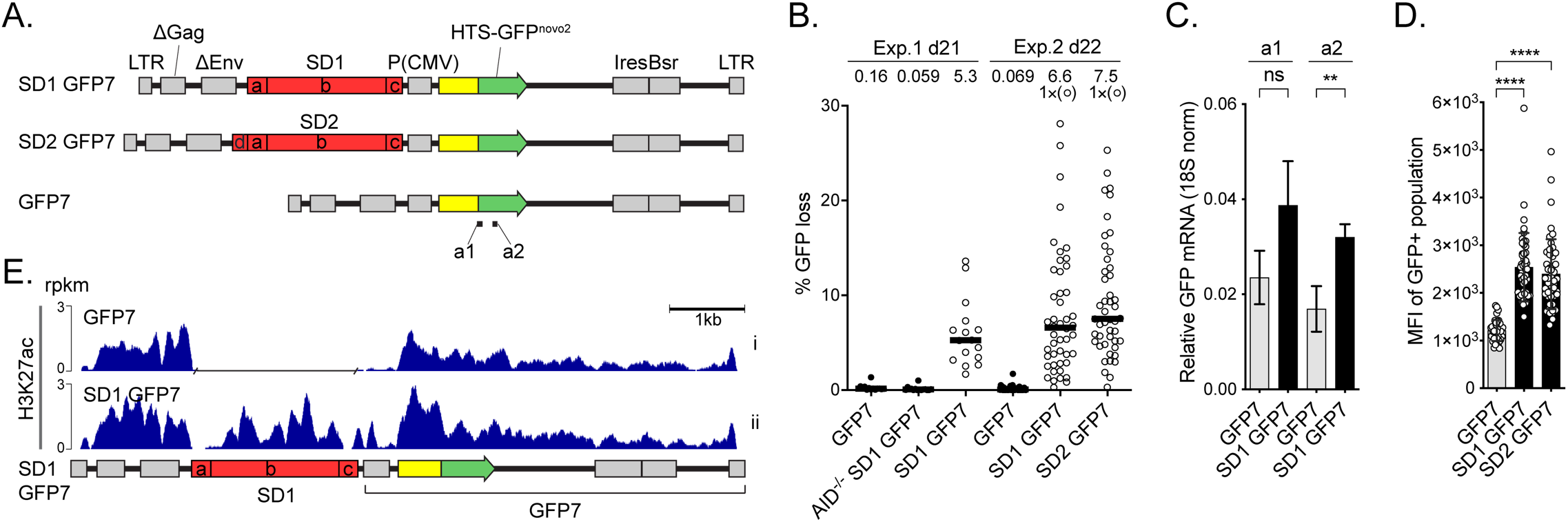
The effect of DIVAC on the H3K27ac in Ramos cells **A**. Outline of the lentiviral *GFP7* reporter used in Ramos cells with SuperDIVAC1 (*SD1 GFP7*), SuperDIVAC2 (*SD2 GFP7*), or without a DIVAC (*GFP7*) inserted upstream of the transcription unit. Locations of amplicons a1 and a2 for RT-qPCR (see C) are indicated. **B**. GFP loss of the *GFP7* reporters in two independent experiments after 21 and 22 days. *SD1 GFP7* was also assayed in AID-deficient Ramos cells (AID^-/-^ SD1 GFP7). Data points outside the y-axis range are in parentheses. Values are medians. **C**. Expression of GFP mRNA from integrated *SD1 GFP7* reporter assessed using two different RT-qPCR amplicons (a1 and a2) indicated in A. Values are mean +/- SD. Student’s t-test * *p*<0.05, ** *p*<0.01, *** *p*<0.001, **** *p*<0.0001, ns not significant. **D**. Mean fluorescence intensity (MFI) of GFP-positive populations of the reporters in Ramos cells. **E**. H3K27ac ChIP-seq of the lentiviral *GFP7* reporter with SD1 (*SD1 GFP7*) and without a DIVAC element (*GFP7*) in Ramos cells. A gap has been inserted in the track of *GFP7* reporter in the place of the SD1 sequence after the alignment. Values are rpkm.

ChIP-seq analysis of H3K27ac in Ramos cells harboring either *GFP7* or *SD1 GFP7* demonstrated that SD1 is associated with H3K27ac, and that the DIVAC induced a modest increase of H3K27ac in the portion of the vector most proximal to the DIVAC (Figure 3E). This increase of signal (Figure 3E) might reflect a spread of H3K27ac writer activity recruited by SD1 into its flanking regions.

Together with H3K27ac and transcription, trimethylation of histone H3 lysine 36 (H3K36me3), a modification associated with bodies of actively transcribed genes with various transcription-associated roles, has predictive power for AID targeting (Li et al., 2019; Wang et al., 2014a). H3K36me3 levels at the transcription unit were elevated by DIVAC 2-3, most prominently downstream of *GFP* (Figure 2A, panels iii and iv), suggesting that DIVAC can modulate transcription and/or co-transcriptional events. H3.3 is an ssDNA-stabilizing histone 3 variant that is enriched in AID target genes (Aida et al., 2013; Romanello et al., 2016). DIVAC 2-3 had no discernable effect on H3.3 levels in the *GFP* open reading frame, arguing that DIVACs do not function by modulating H3.3 in the SHM-targeted region (Figure 2A, panels v and vi). While these epigenetic features of the associated genomic region might be needed for SHM, the inherent mutation enhancer activity of the DIVACs does not rely on the modulation of H3K27ac or H3.3.

### Neither antisense nor convergent transcription are required for efficient SHM

To assess the effect of DIVAC on antisense transcription at the GFP reporter, we applied global run-on sequencing (GRO-seq) to subclones of DT40 cells with the *Ri GFP2* and *Ri GFP2 2-3* reporters. This assay measures the location and abundance of elongation-competent Pol2 (Core et al., 2008). There were no detectable changes in sense transcription of the *GFP* gene, antisense transcription upstream of the promoter or around the polyA-signal sequence between the two reporters (Figure 4A). We did not observe any significant antisense transcription in the region of the reporter where most of the mutations occur (Figure 4A and see below). Thus, we found no evidence for convergent or antisense transcription in the areas targeted for SHM.

**Figure 4.**
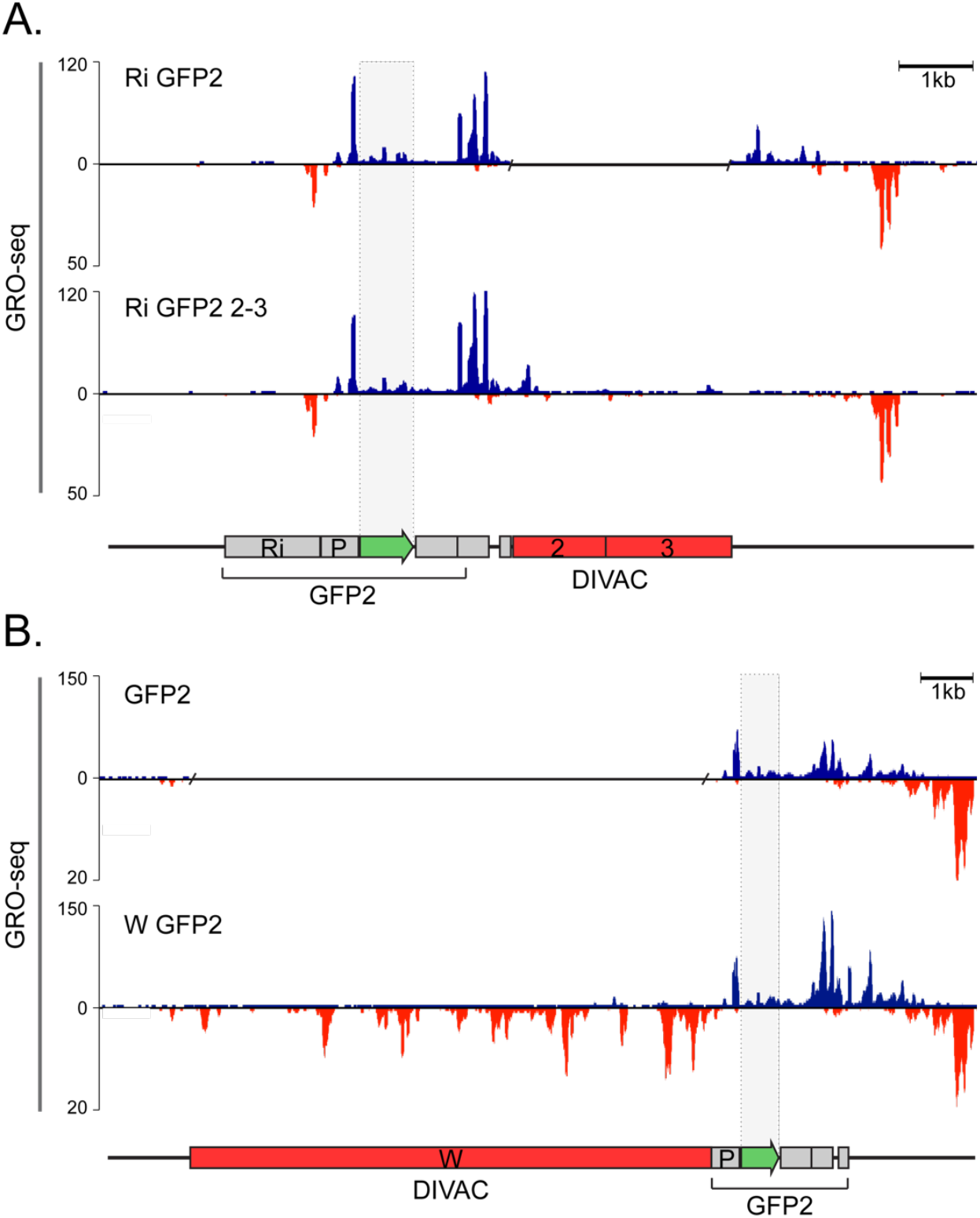
DIVACs have no effect on antisense transcription **A**. GRO-seq analysis of the genome-integrated *Ri GFP2* (upper panel) and *Ri GFP2 2-3* reporters (lower panel). **B**. GRO-seq analysis of the *GFP2* (upper panel) and *W GFP2* (lower panel) reporters. The upper track of each panel represents the sense transcription (blue) and the lower panel the antisense transcription (red). Note the differences in the scale. A gap has been inserted in the tracks of reporters that do not have a DIVAC. The region of *GFP* coding sequence has been highlighted with shading. The y axes indicate the GRO-seq counts normalized to number of reads per million.

We also performed GRO-seq with a *GFP2* reporter without a DIVAC (*GFP2*) and with the W DIVAC located upstream of the transcription unit (*W GFP2*). Similar to DIVAC 2-3, W DIVAC did not induce detectable antisense transcription in *GFP*, nor did it detectably increase elongation-competent Pol2 engaged in sense transcription in *GFP* (Figure 4B). This strengthens the conclusion that convergent and antisense transcription in the mutation target region are not required for SHM. We did observe a DIVAC-dependent increase in Pol2 sense transcription at the end of the transcription unit (Figure 4). This increase is present regardless of the location of DIVAC upstream or downstream of the transcription unit and is greater with the stronger W DIVAC (Figure 4B).

We reasoned that if promoter-upstream antisense transcription recruits exosome and underlies the targeting of SHM to the IgV region (Pefanis et al., 2014), we should detect DIVAC-dependent divergent transcription upstream of the promoter of the *GFP2* reporter. The divergent transcription was clearly visible in both the *Ri GFP2* and *Ri GFP2 2-3* reporters, but DIVAC did not increase its magnitude (Figure 4A). No meaningful comparison of upstream divergent transcription between the *GFP2* and *W GFP2* reporters could be performed due to the different DNA sequence immediately upstream of the promoter (Figure 4B).

These data indicate that DIVAC does not lead to or consistently increase divergent or convergent transcription and thus dissociate the production of antisense RNA from the function of DIVACs. Therefore, induction of antisense transcription at the target gene does not play a major role in targeting of mutations by immunoglobulin enhancers in this model system for SHM.

### DIVACs regulate Pol2 progression through the mutating target gene

To investigate whether DIVAC induces mutations by modulating Pol2 progression, we characterized the effect of DIVAC on Spt5 recruitment as well as the phosphorylation status of the Pol2-CTD by ChIP-seq. Visual comparison of the data for the *Ri GFP2 2-3* and *Ri GFP2* reporters suggested that DIVAC did not strongly influence the association of the Pol2 catalytic subunit RPB1 (Figure 5A, tracks i and ii), Spt5 (tracks iii and iv), S5P-CTD (tracks v and vi), or S2P-CTD (tracks vii and viii) in the vicinity of the TSS. To examine this quantitatively, we calculated the ratio of the ChIP signal in the promoter-proximal region (TSS ±500 bp, Figure 5B) of the *GFP* gene for the Ri GFP2 2-3 versus the control Ri GFP2 cell lines and compared that ratio to a similar ratio calculated for other highly transcribed genes in the genome, which should not be affected by the presence or absence of DIVAC in the reporter. We found that this ratio was not increased at the *GFP* gene relative to other well-expressed genes (see Materials and Methods) for any of the four features assessed by ChIP-seq (Figure 5C, bars labeled “Prom”). This indicates that DIVAC does not increase promoter-proximal levels of total RPB1, its phosphorylated forms, or Spt5 in DT40 cells, and argues that DIVAC does not regulate promoter-proximal pausing.

**Figure 5.**
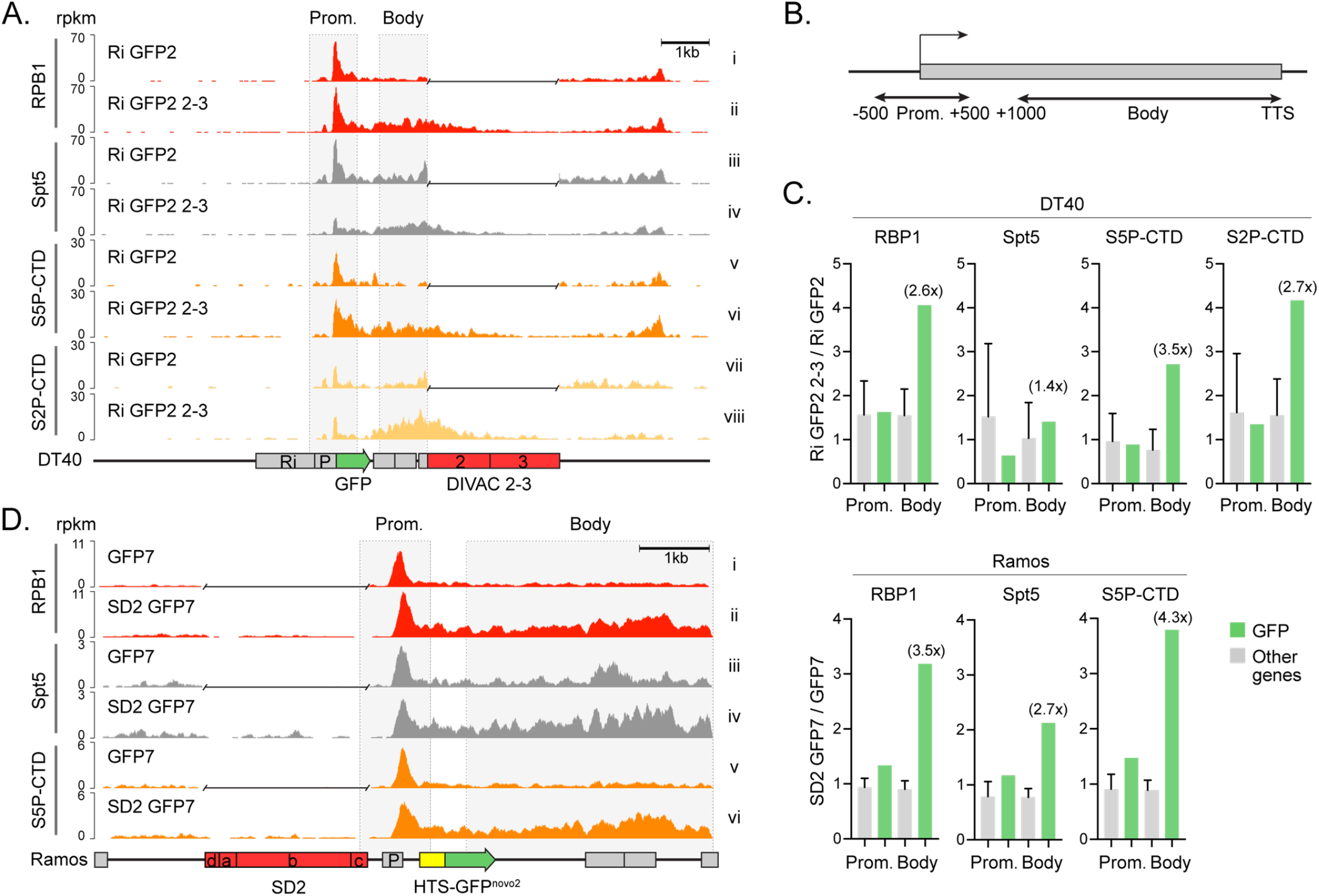
DIVACs induce polymerase stalling **A**. ChIP-seq analysis of the RPB1, Spt5, S5P-CTD, and S2P-CTD of the genome-integrated *Ri GFP2* (upper tracks) and *Ri GFP2 2-3* (lower tracks) reporters and their surrounding region in DT40 cells as indicated. **B**. The regions for the promoters (TSS +/-500 bp) and the bodies (TSS +1,000 bp to TTS) of genes analyzed in C. **C**. Ratios of ChIP-seq signal (rpkm) in DT40 and Ramos cells with and without a DIVAC in the regions indicated in B. The ratios at the *GFP* transcription unit (green bars) were compared to the ratios of 500 most highly expressed genes (other genes, gray bars). The fold change at the GFP compared to other genes in the gene body is indicated in parentheses. The data are mean + SD. **D**. ChIP-seq analysis of the RPB1, Spt5, and S5P-CTD of the *GFP7* (upper tracks, no DIVAC) and *SD2 GFP2* (lower tracks) reporters in the Ramos cells as indicated. A gap has been inserted in the tracks of reporters without a DIVAC after the alignment.

The ChIP-seq traces instead indicated DIVAC-dependent increases in all four parameters in the body of the transcription unit, particularly for S2P-CTD (Figure 5A). Repeating the quantitative analysis of the ChIP-seq signals as before, but now in gene bodies (defined as 1,000 bp downstream of the TSS to the transcription termination signal, TTS, Figure 5B), revealed clear DIVAC-dependent 2.6- to 3.5-fold increases of total Pol2, S5P-CTD, and S2P-CTD in the *GFP2* gene body in comparison to other highly expressed genes in DT40 cells (Figure 5C, bars labeled “Body”). Despite the visually apparent DIVAC-dependent increase of Spt5 in the *GFP2* gene body (Figure 5A, tracks iii and iv), the quantitated increase was less striking than for the other parameters (Figure 5C). These data argue that DIVAC induces the accumulation of Pol2, S5P-CTD, and S2P-CTD in the gene body in a context in which there is no increase in the production of RNA transcripts or GFP protein and no increase in the accumulation of elongation-competent Pol2. We note that the region where the DIVAC-dependent increase in H3K36me3 signal was observed (Figure 2A) is encompassed by the region with DIVAC-dependent increases in Pol2 and its CTD-phosphorylated forms, consistent with the possibility of S2P- and S5P-CTD-mediated recruitment of H3K36me3 writer activity in the DIVAC-regulated transcription unit (Figure 5A, tracks v to viii).

We then tested whether DIVAC has similar effect in avian and human cell lines and in a different reporter configuration, in particular, moving DIVAC to a position upstream of the transcription unit. RPB1, Spt5, and S5P-CTD ChIP-seq analyses of Ramos cells transduced with either the *GFP7* or *SD2 GFP7* reporters revealed increases in these factors in the transcription unit (Figure 5D). Quantitation of the ChIP-seq signals confirmed strong DIVAC-dependent increases in RPB1, S5P-CTD, and to a somewhat lesser extent Spt5 in the body of the *GFP7* transcription unit compared to other highly expressed genes (Figure 5C). These increases in the gene body were much larger than in the vicinity of the promoter, recapitulating the observation from DT40 data (Figure 5C). Therefore, DIVAC-induced Pol2 accumulation is not an idiosyncrasy of a particular cell line or configuration of the reporter system, and the phenomenon is conserved across species.

Taken together, these findings demonstrate that DIVACs induce Pol2 stalling in the bodies of SHM target genes and, importantly, that these increases are accompanied by little or no increase in the production of mature mRNA.

### Single-stranded DNA regions can be increased by a DIVAC but do not predict mutability

Some evidence suggests that IgV regions and perhaps also non-Ig targets of AID have substantial ssDNA character, which is presumed to promote the ability of AID to act on those regions (Parsa et al., 2012; Romanello et al., 2016; Ronai et al., 2007). However, it is unclear whether ssDNA is sufficient prerequisite for SHM, or whether susceptibility to the action of AID requires ssDNA in a specific context, in which case many or even most ssDNA regions might not be efficient targets of AID. The capacity of DIVAC to stall polymerase implies increased single-strandedness in DIVAC-regulated genes, but the effect of DIVAC on ssDNA in unknown.

To assess the contribution of ssDNA patches to the locus-specificity of SHM and to test the hypothesis that DIVAC increases ssDNA character in SHM target regions, we analyzed the effect of various DIVACs on the amount of ssDNA in the *GFP* transcription unit using an established *in situ* bisulfite assay (Ronai et al., 2007). We treated cross-linked nuclei with bisulfite to sulfonate accessible single-stranded deoxycytidines (ssCs) in the chromatinized genome, and after deproteinization, identified the bisulfite-converted ssCs in the *GFP2* transcription unit by sequencing. From the sequencing data, we calculated the frequency of ssCs (referred to hereafter as ssDNA frequency) and ssDNA patch size (see Material and Methods). To single out AID-induced mutations, the assay was performed in parallel in AID-deficient cells (*AID*^*-/-*^ *W GFP2*). *AID*^*-/-*^ *W GFP2* had a similar ssDNA frequency as the AID expressing cells (10 × 10^−3^ and 9.5 × 10^−3^, respectively, *p*=0.76, Figure 6A), indicating that AID expression does not bias the assay readout and that DIVAC can increase single-strandedness in AID-independent manner. As expected, we found no ssDNA patches in a non-transcribed control gene *keratin 5* (ssDNA frequency <1.4 × 10^−3^, 4,049 bp sequenced).

**Figure 6.**
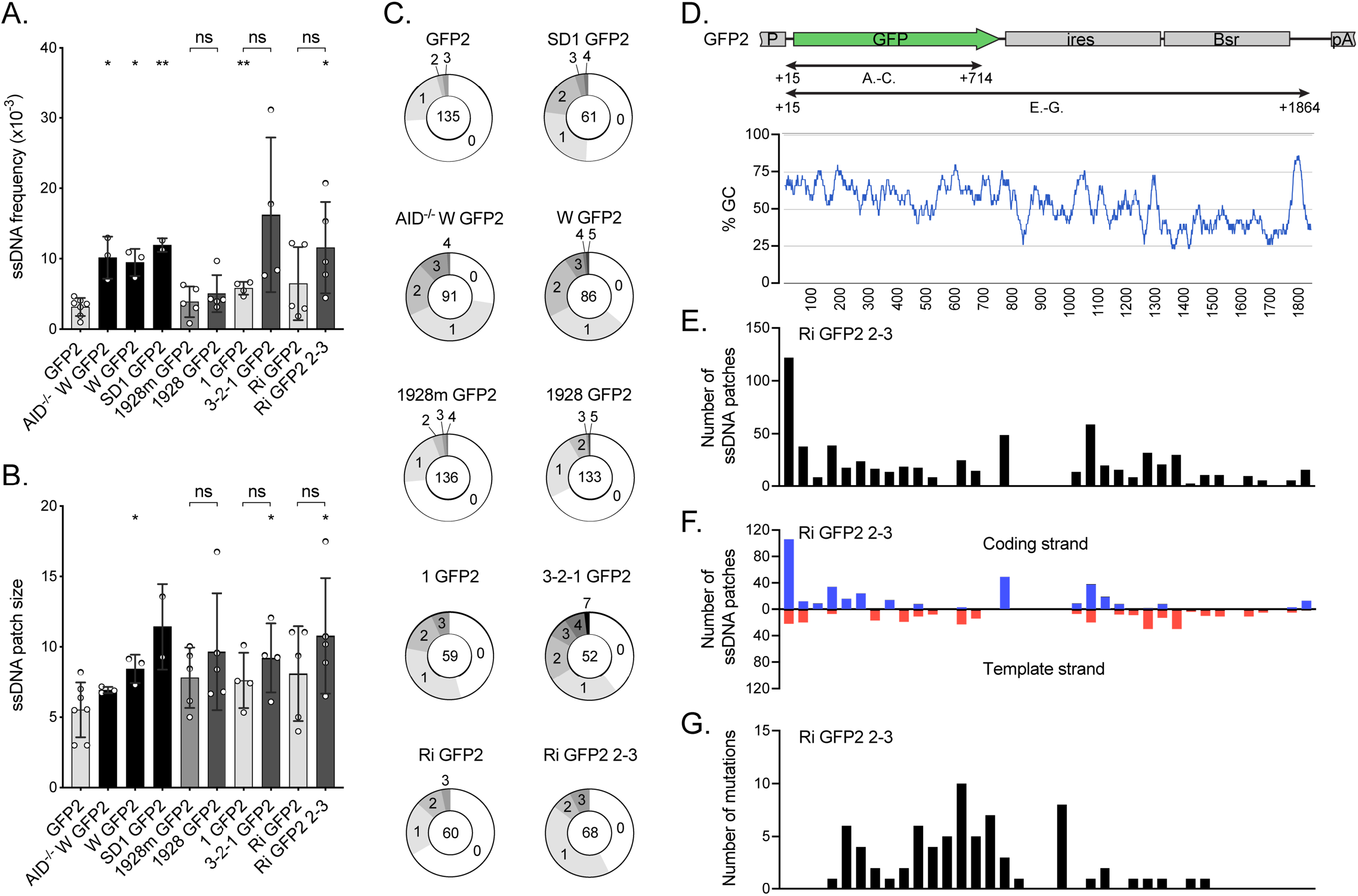
The effect of DIVACs on ssDNA **A**. The frequency of ssDNA in the reporters assayed by *in situ* bisulfite assay (ratio of modified ssCs in patches to all the sequenced Cs) in the region TSS +15 bp to TSS +714 bp as indicated in D. To be counted as a single-stranded patch, a minimum of 2 consecutive C-to-T or G-to-A mutations were required. The asterisks indicate statistical significance according to Welch’s t-test when compared to *GFP2*. Lack of significance of additional comparisons are indicated. * *p*<0.05, ** *p*<0.01, ns not significant. The shading of the bars is as in Figure 1B. **B**. The size of ssDNA patches (mean of minimum and maximum possible patch size) in the same region of the reporters as in A. The asterisks indicate statistical significance according to Welch’s t-test when compared to *GFP2*. Lack of significance of additional comparisons are indicated. * *p*<0.05, ns not significant. The shading of the bars is as in Figure 1B. **C**. Distribution of ssDNA patches detected per sequence in the same region of the reporters as in A. The numbers indicate the numbers of patches per sequence and the number in the middle indicates the total number of analyzed sequences. **D**. The location (upper panel) and GC content (bottom panel) of the regions along the *GFP2* transcription unit in which the ssDNA and mutations were analyzed in A-C as well as in E-G. The GC content is plotted with a 30-bp window. **E**. Location of ssDNA patches (center) in *Ri GFP2 2-3* reporter transcription unit (TSS +15 bp to TSS +1,864 bp). The location of ssDNA patches is plotted in bins of 50 bp. **F**. The location of ssDNA patches on the coding strand (blue, above the x-axis) and template strand (red, below the x-axis) of the *Ri GFP2 2-3* reporter. The locations are plotted in bins of 50 bp. **G**. The number of mutations located along the Ri GFP2 2-3 reporter are plotted in bins of 50 bp. The graphs in E-G align with D.

The bisulfite sequencing data revealed that ssDNA frequency at the *GFP* coding sequence was generally higher in the presence of a functional DIVAC than in its absence. Reporters lacking an active DIVAC had ssDNA frequencies ranging from 3.2 × 10^−3^ for *GFP2* to 6.5 × 10^−3^ for *Ri GFP2* (Figure 6A). A statistically significant ∼3-fold increase in ssDNA frequency was seen with the DIVACs W (9.5 × 10^−3^, *p*=0.016) and SD1 (12 × 10^−3^, *p*=0.0066) compared to *GFP2* (Figure 6A). The highest ssDNA frequencies were observed with the *3-2-1 GFP2* (16 × 10^−3^), *SD1 GFP2* and *Ri GFP2 2-3* (12 × 10^−3^) reporters, representing a significant increase over *GFP2* in the case of *Ri GFP 2-3* (*p*=0.043) and a 2.8-fold increase in the upstream position and 1.8-fold increase in the downstream position relative to their respective control vectors *1 GFP2* (5.8 × 10^−3^, *p*=0.15) and *Ri GFP2* (6.5 × 10^−3^, *p*=0.21), although these differences were not statistically significant due to large variations in the datasets (Figure 6A). DIVAC 1928 represents the one exception to the correlation of ssDNA frequency with DIVAC activity. While it induces SHM more than 100-fold over the *GFP2* control (Figure 1B), the ssDNA frequency in *1928 GFP2* (5.0 × 10^−3^) is smaller than in some reporters lacking a DIVAC (e.g., *Ri GFP2* and *1 GFP2*) and is only 1.6-fold greater than in *GFP2* (*p*=0.19) (Figure 6A). Furthermore, mutation of its E boxes had a minimal effect on ssDNA frequency (*1928m GFP2*, Figure 6A) while reducing DIVAC activity 11-fold (Figure 1B). Together, these findings indicate that ssDNA frequency at SHM target regions detected by bisulfite accessibility exhibits a correlation with but does not reliably predict DIVAC activity. The difficulty in establishing a tighter correlation might stem from the effects of upstream DIVAC elements on transcription efficiency (as reflected by GFP MFI; Figure 1C) and limitations in the ability of the bisulfite assay to specifically detect ssDNA regions relevant to SHM (see below and Discussion).

Intriguingly, ssDNA patch size was consistently larger for reporters containing a strong DIVAC element than those without DIVAC function, ranging from 5.5 nt in *GFP2* to 11 nt in *SD1 GFP2* and *Ri GFP2 2-3* (Figure 6B). *W GFP2* (8.4, *p*=0.016), *3-2-1 GFP2* (9.2, *p*=0.047), and *Ri GFP2 2-3* (11, *p*=0.042) had significantly larger ssDNA patches than the *GFP2*. However, these differences were not statistically significant when *3-2-1 GFP2* and *Ri GFP2 2-3* were compared to their respective controls *1 GFP2* and *Ri GFP2*.

The distribution of ssDNA patches across the analyzed sequences reflects the frequency of ssDNA patches per sequence. The strongest DIVACs induced more patches (W and 1928 up to 5 and 3-2-1 up to 7) per sequence than weaker DIVACs or reporters without a DIVAC, where most sequences did not have any patches (Figure 6C).

Analysis of the distribution of ssDNA patches along the *GFP2* transcription unit (TSS +15 to +1,864 bp) in *Ri GFP2 2-3* revealed that the location of ssDNA patches did not correlate well with the location of mutations or GC distribution (Figure 6D-G). Apart from a peak immediately downstream of the TSS, ssDNA patches were fairly evenly distributed along the *GFP2* transcription unit (Figure 6E) and did not display an obvious strand bias (Figure 6F). A similar distribution of ssDNA patches was seen with upstream DIVAC in the reporter *3-2-1 GFP2* (Figure S1). Mutations, however, accumulated largely in the region between TSS +200 and TSS +1,000 bp (Figure 6G), a pattern reminiscent of that observed during SHM of Ig genes (Odegard and Schatz, 2006). We conclude that the bulk of the single-stranded DNA detected by the *in-situ* bisulfite assay in the mutating gene is unlikely to be a direct consequence of SHM-enhancing activity of DIVACs and that SHM is directed to the TSS +200 to +1,000 bp region via a mechanism that is not dictated by the incidence of ssDNA alone.

Together, our data argue that DIVACs mediate their function by stalling the elongating RNA Pol2 complex within the SHM target gene. This could allow AID that travels with the Pol2 along with its elongation complex DSIF-PAF-SPT6 to take advantage of the availability of increased ssDNA, stabilized by the stalled Pol2.

## DISCUSSION

Pol2 stalling has been linked to CSR through several findings. The polymerase accumulates when entering GC-rich switch regions (Rajagopal et al., 2009; Wang et al., 2009), whose GC-bias contributes to a potential to form unusual DNA structures such as R-loops and G-quadruplexes that might affect polymerase progression (Yeap and Meng, 2019; Yu and Lieber, 2019). While Pol2 stalling has been suggested to be involved in the targeting of SHM (Canugovi et al., 2009; Maul et al., 2014; Wang et al., 2014b), the connection has remained tenuous, largely due to a lack of explanation for how Ig locus-specificity might be achieved. Our findings show that Ig enhancers increase Pol2 stalling in the SHM target region, providing a plausible explanation for how they accomplish SHM targeting.

Transcription is required for SHM, likely for several reasons: 1) transcription is thought to provide the ssDNA template for AID (Casellas et al., 2016); 2) AID associates with Pol2 and Pol2-associated factors such as PAF1, Spt5, Spt6, and Cdk9 (Begum et al., 2012; Methot et al., 2018; Pavri et al., 2010; Willmann et al., 2012); 3) transcription provides RNA that can stabilize AID association with DNA (Qiao et al., 2017), target AID to the target region either directly (Zheng et al., 2015) or via RNA processing and splicing factors such as RNA exosome (Basu et al., 2011) and SRSF1-3 (Kanehiro et al., 2012; Singh et al., 2020) among others; and 4) transcription might help maintain accessible DNA and appropriate epigenetic marks. However, neither transcription nor AID association with chromatin predicts mutability (Feng et al., 2020). Instead, the specificity of SHM for Ig loci appears to be dictated by Ig enhancers and enhancer-like elements (DIVACs) by an unknown mechanism (Blagodatski et al., 2009; Buerstedde et al., 2014). In the context of Ig loci, it is likely that DIVACs have two distinct SHM-promoting functions, as transcriptional enhancers and as mutational enhancers. While these activities might usually be coupled, they can be physically separated and perhaps rely on binding of different factors (Buerstedde et al., 2014; Kothapalli et al., 2011). However, almost nothing is known about how DIVACs, as mutation enhancers, might alter the properties of the target gene.

To identify features specifically associated with the mutation enhancer activities of DIVAC, we took advantage of a system in which target gene expression is largely or completely independent of DIVAC but in which DIVAC can stimulate SHM by two orders of magnitude. Using this system, we investigated multiple target gene features previously implicated in SHM targeting.

We found that several of these features are either not required for robust SHM targeting or not regulated by the mutation enhancer activity of DIVACs. First, while convergent transcription appears to explain some off-target activity of AID (Meng et al., 2014), our data demonstrate that it is not stimulated by DIVAC or required within the target region for mutations to occur, at least in the context of our SHM reporters. Indeed, our findings demonstrate that robust SHM can occur in the absence of substantial antisense transcription. In addition, we find that promoter-upstream antisense transcription is not affected by DIVAC, suggesting that RNA exosome activity, previously suggested to play a role in AID IgV targeting and off-targeting (Pefanis and Basu, 2015), is not a major target of regulation by mutation enhancers. Second, H3K27ac, a marker used to define super-enhancers, is associated at approximately equal levels with the mutation target region in the presence and absence of mutation enhancers. DIVAC itself is abundantly marked by H3K27ac, and this additional H3K27ac in the vicinity of the SHM target region could increase the mutability of the target region. Notably, a 10-fold reduction of mutation enhancer activity, achieved by mutating the E boxes in a DIVAC, did not reduce H3K27ac levels. Third, the histone variant H3.3, which has the potential to stabilize AID-accessible ssDNA regions in the chromatin of DT40 cells (Romanello et al., 2016), was not strongly induced by DIVACs. Our data do not address the possibility that H3K27ac, H3.3, or promoter-upstream antisense RNA production are needed for SHM, as they are present with and without DIVACs, but our findings strongly argue that they are not sufficient to induce SHM, are not the primary mechanism by which mutation enhancers operate, and, therefore, also not likely to be central to the mechanism that confers Ig-locus specificity to SHM.

Our data show that DIVAC increases the presence of Pol2, Spt5, S5P-CTD, and S2P-CTD at the mutating transcription unit. In striking contrast, the gene product (GFP) and transcription at the transcription unit (run-on) were not increased, and mature mRNA production was not or only slightly increased. The lack of increased transcriptional output argues that DIVAC prevents the progression of some Pol2 within the transcription unit. This could be achieved by mechanisms that include induced stalling, backtracking, or premature termination of Pol2.

As DIVAC does not reduce gene output, the apparent reduction in Pol2 progression might be compensated for by an increase of Pol2 loading, reinitiation, and/or elongation complex formation. Our data indicate that mutation enhancers support robust S2P-CTD modification of Pol2, likely by P-TEFb, consistent with substantial pause release, suggesting that the mechanism by which DIVAC stalls Pol2 is not similar to that of promoter-proximal pausing. Thus, it is possible that DIVAC impedes Pol2 progression via an external stalling force such as by DNA topological stress. To release the stalling, Pol2 backtracking could be involved and the exposed 3′ end of RNA subjected to degradation by TFIIS or RNA exosome complex, leading to ssDNA exposure and subsequent cytidine deamination. Indeed, stalling of Pol2 by a premature transcription termination signal or excessive positive supercoiling ahead of the elongating Pol2 can increase the amount of Pol2, AID, and the frequency of SHM (Kodgire et al., 2013; Maul et al., 2015).

DIVAC also increased levels of the histone mark H3K36me3 in the transcription unit. Histone methyltransferase SETD2 selectively binds S5P-S2P dually phosphorylated Pol2 during transcription elongation (Kizer et al., 2005; Li et al., 2005; Vojnic et al., 2006) and is the only known writer of H3K36me3 (Edmunds et al., 2008). While our analyses do not assess dually phosphorylated CTD, it is possible that DIVAC often induces S2P and S5P in the same CTD. H3K36me3 has various important functions including preventing transcription initiation in gene bodies by, e.g., recruiting histone chaperones such as FACT complex (Carvalho et al., 2013), facilitating DNA repair (Sun et al., 2020), and regulating RNA splicing (Barash et al., 2010). Several protein factors mediating these biological processes have been directly or indirectly implicated in AID recruitment, mutation targeting, or SHM activity (Aida et al., 2013; Kanehiro et al., 2012; Methot et al., 2018) and might be involved in reinforcing the restriction of SHM within a DIVAC-containing TAD (Senigl et al., 2019).

In a striking parallel with DIVAC-mediated regulation of its target gene, SHM-susceptible or “hot” TADs genome-wide are significantly enriched in Spt5, and S5P-CTD, but not with increased GRO-seq or H3K4me3 signal when compared to SHM-resistant or “cold” TADs (Senigl et al., 2019). Furthermore, at least some hot TADs contain elements with DIVAC activity (Senigl et al., 2019). Combined with these observations, our findings raise the possibility that on-target and off-target SHM have mechanistic similarities.

Our findings suggest that DIVAC can increase the single-strandedness of the mutating gene. The DIVAC-induced patches roughly correspond to the size of the Pol2 transcription bubble and the size of AID-induced clustered mutations during Pol2 transcription *in vitro* (Parsa et al., 2012; Pham et al., 2019; Romanello et al., 2016; Ronai et al., 2007). One potentially confounding variable in our analysis of ssDNA frequency is overall transcription levels. The strongest DIVACs that induce ssDNA character (W, SD1, and 3-2-1) were positioned upstream of the transcription unit and increased the expression of the GFP most strongly, likely due to their proximity to the promoter. Interestingly, the distribution of ssDNA patches does not resemble the mutation pattern and instead resembles the distribution of Pol2 and Spt5 in the reporter (Figure S2). Thus, ssDNA patches detected with the *in-situ* bisulfite method are likely to track Pol2 complexes, including the vast majority that do not lead to AID-mediated deamination. Consistent with a contribution of transcription to ssDNA frequency, mutation of the E boxes in DIVAC 1928 has little effect on either GFP MFI, transcription (McDonald et al., 2013), or ssDNA frequency.

Transcription bubbles created by the Pol2 complex cannot by themselves be what targets SHM. The bubbles are accessible to nucleotides but not proteins (Barnes et al., 2015), and the reporter without a DIVAC is robustly transcribed but not mutated, as are many genes in GC B cells, including Ig constant regions. Even in rearranged IgV regions of GC B cells, where SHM is occurring efficiently, only approximately one mutation occurs per cell division, highlighting the fact that very few Pol2 transcription events lead to a mutation (Yeap and Meng, 2019). Therefore, it is likely that only a small subset of Pol2 complexes in the mutation target region are competent to recruit AID action and SHM. If indeed only a small fraction of Pol2 is relevant for SHM, identification of those Pol2 molecules might be difficult.

Importantly, DIVAC 2-3 in the downstream position (*Ri GFP2 2-3*) does not increase GFP expression (at either protein or mRNA level) but increases the ssDNA character of the target. This indicates that in this case, DIVAC-induced ssDNA is not simply a result of increased transcriptional output. We found that the size of ssDNA patches was consistently increased in the presence of an active DIVAC. Such an increase could arise in a local region of negative supercoiling that stabilizes ssDNA, e.g., upstream of clustered, arrested Pol2-complexes (Wang et al., 2014b). Indeed, bisulfite-accessible ssDNA can arise from DNA negative supercoiling that facilitates AID access to DNA (Parsa et al., 2012). Together, our findings suggest that DIVAC can increase the single-strandedness of the mutating gene and that the induced single-strandedness could be linked to the mutation enhancer activity of DIVAC.

In the absence of histone H3.3, ssDNA, SHM, and GCV are reduced in the *IgL* V region of DT40 cells without affecting transcription kinetics, suggesting that H3.3 can promote or stabilize ssDNA and thereby facilitate IgV region mutation (Romanello et al., 2016). H3.3 is also associated with many AID off-targets genome-wide (Aida et al., 2013). However, DIVAC does not have a major effect on levels of H3.3 at the *GFP* gene. Thus, while H3.3 can facilitate AID-mediated diversification, it is unlikely to be critical for determining the locus-specific targeting of SHM.

The catalytic pocket of AID is shielded and AID catalysis is remarkably slow, mediating a reaction every 1–4 minutes (King et al., 2015; Larijani et al., 2007), and slow catalysis is potentially an important rate-limiting step for SHM. Longer persistence of AID on DNA with stalled Pol2 could allow sufficient time for AID to act on ssCs. We cannot rule out the role of AID recruitment by DIVACs, as AID ChIP signals are not consistently above background even in the context of a strong DIVAC (data not shown). Given that Pol2-associated Spt5 can recruit AID to DNA (Maul et al., 2014; Methot et al., 2018; Pavri et al., 2010), even a small local increase of Spt5 and Pol2 by DIVAC, particularly in the context of stalled Pol2, might have a significant impact on the efficiency of SHM.

In summary, our data implicate Pol2 elongation control in the function of DIVACs and the targeting for SHM and support a model in which Ig enhancers allow more time for AID to act on ssDNA in the SHM target region, thereby increasing the likelihood of a deamination event.

## Acknowledgements

The work was supported by the Israel Science Foundation grant 1920/20 (Y.M.), Czech Science Foundation grant 15-24776S (F.S.), National Institutes of Health grants AI 127642 (D.G.S.) and T32 AI 007019 (J.J.M.), grants from the Sigrid Juselius Foundation, the Jane and Aatos Erkko Foundation, the Jenny and Antti Wihuri Foundation, the Ella and Georg Ehrnrooth Foundation, the Cancer Society of South-West Finland and the Emil Aaltonen Foundation (J.A.), the Turku University Foundation and the Maud Kuistila Memorial Foundation (J.A. and A.S.), as well as K. Albin Johanssons Foundation (P.M.).

---

**Figure S1.**
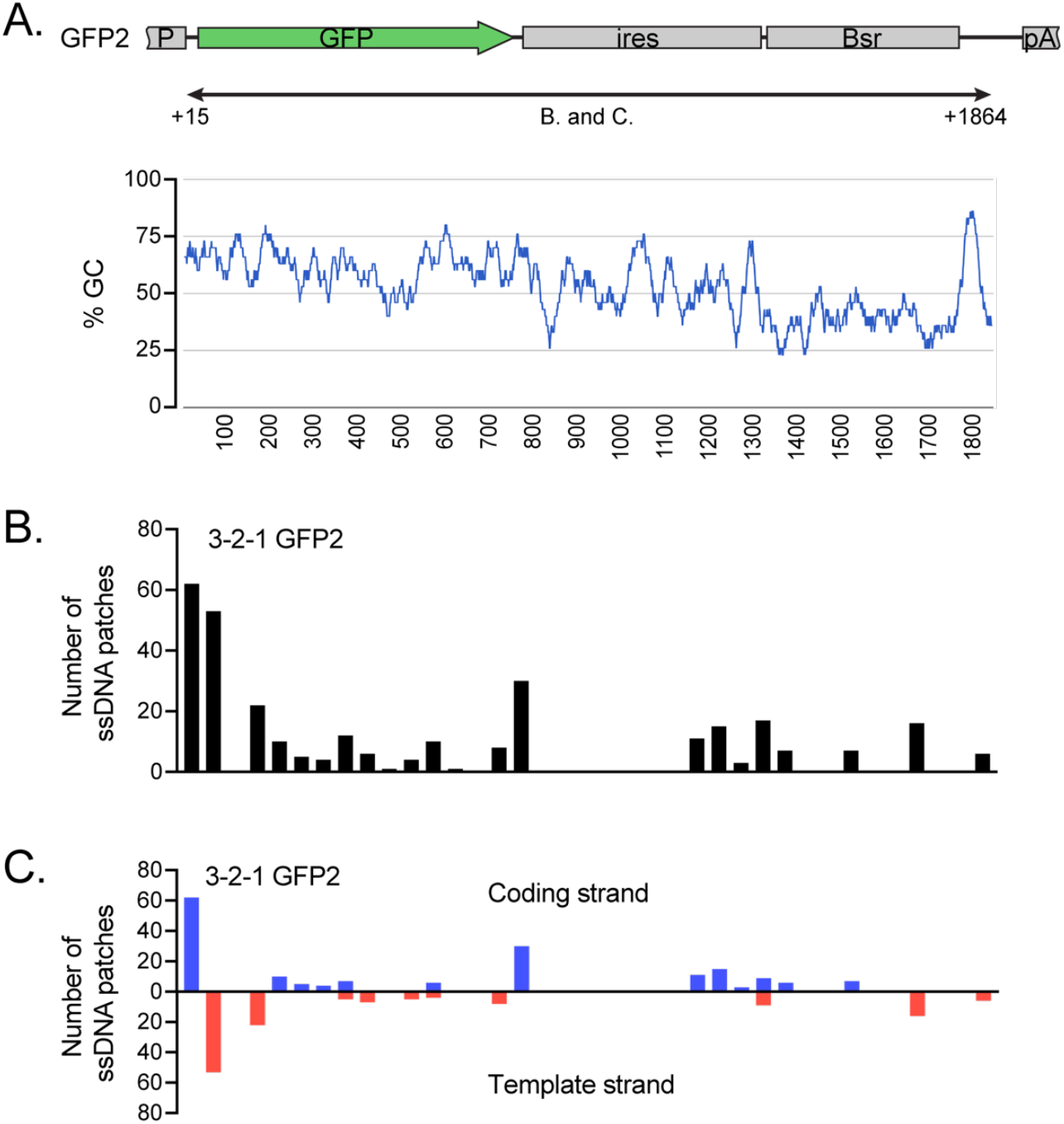
The effect of 3-2-1 DIVAC on ssDNA patch location **A**. The location (upper panel) and GC content (bottom panel) of the region along the *GFP2* transcription unit in which the ssDNA was analyzed for B and C. The GC content is plotted with a 30 bp window. **B**. Location of ssDNA patch centers in *3-2-1 GFP2* reporter plotted in bins of 50 bp. **G**. The location of ssDNA patches on the coding strand (blue, above the x-axis) and template strand (red, below the x-axis) of the *3-2-1 GFP2* reporter. The locations are plotted in bins of 50 bp.

**Figure S2.**
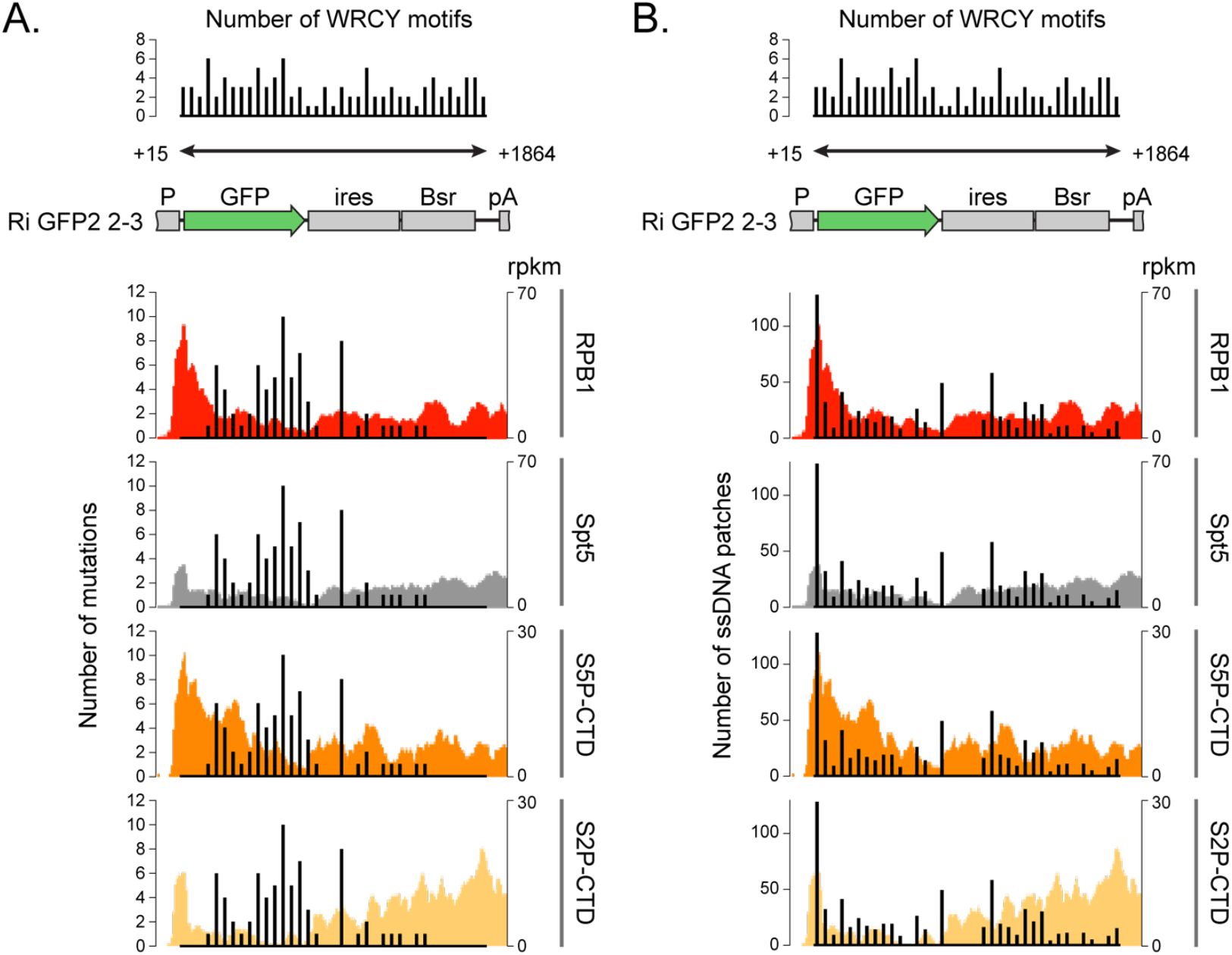
Colocation of Pol2, mutations and ssDNA patches Comparison of the ChIP-seq signal (from Figure 5A, left y axes) with the location of AID-induced mutations (A) and the location of ssDNA patches (B) at the *Ri GFP2 2-3* reporter plotted in bins of 50 bp (y axes). The location of AID hotspot motifs WRCY/RGYW along the sequenced region (TSS +15 bp to +1,864 bp) is shown at the top.

